# Fitting a square beam in a square camera: novel condenser apertures for low-dose transmission electron microscopy

**DOI:** 10.1101/2023.08.13.553155

**Authors:** Hamish G. Brown, Dan Smith, Benjamin C. Wardle, Eric Hanssen

**Affiliations:** Ian Holmes Imaging Centre and ARC Centre for Cryo Electron Microscopy of Membrane Proteins, The Bio21 Molecular Science and Biotechnology Institute, The University of Melbourne, Parkville, Victoria, 3010, Australia; Melbourne Centre for Nanofabrication, Australian National Fabrication Facility, Clayton, VIC 3168, Australia; Materials Characterisation and Fabrication Platform (MCFP), Australian National Fabrication Facility – Melbourne Node, The University of Melbourne, Parkville, Victoria, 3010, Australia; Thermo Fisher Scientific Australia, 5 Caribbean Dr, Scoresby VIC 3179, Australia; Department of Biochemistry and Pharmacology, The University of Melbourne, Parkville, Victoria, 3010, Australia

## Abstract

In transmission electron microscopy (TEM) cameras are square or rectangular but beams are round. With a beam size chosen to fill the camera at a given image magnification, the circular lobes of the beam will extend beyond the camera’s field of view and irradiate areas that are not acquired on the camera, damaging and precluding them from future acquisitions if the sample is beam sensitive. In this paper we present development of condenser aperture plates for TEM that have square and rectangular apertures which improve the efficiency of sample area usage by 44% or greater in low dose TEM applications. We demonstrate that the use of these apertures is compatible with high-resolution cryogenic (cryo) TEM by reconstructing sub 2 Å apo-ferritin models from a datasets recorded with both square and rectangular apertures. Moreover the design of our aperture plates should improve the flexibility of 2 condenser systems for cryo-TEM acquisitions with multiple shots per hole by tailored matching of beam sizes to camera sizes at each magnification.

## INTRODUCTION

The past decade has seen what some observers have dubbed a ‘resolution revolution’ in the single particle analysis (SPA) technique in cryogenic transmission electron microscopy (cryo-TEM) [1]. In this technique tens to hundreds of thousands of single particles embedded in vitreous (amorphous) ice are imaged in the TEM at high magnification. Biological material is beam sensitive and only permits a dose typically around 40 e/Å^2^ before even low resolution information about the molecule is lost [2]. With this low a dose, even under ideal imaging conditions single images of intact protein molecules do not exhibit high resolution due to low signal-to-noise. Sophisticated software can combine the thousands of low dose images of the particle in different orientations into a high resolution 3D map [3]. As more quality images of particles are added to the dataset, the reconstructed map typically improves as the reconstruction algorithm can assign better confidence to weak high-resolution signals [4], thus the number of images that can be routinely recorded in a given amount of time and on a single cryo-EM grid is important for the technique.

The aforementioned resolution revolution has seen recent exponential growth in the number of structures solved using this technique, and the typical resolutions of these maps have also drastically improved [1]. The SPA technique has been demonstrated at near atomic resolution, 1.22 Å [5], with the average resolution of a map submitted to the protein database (PDB) being just below 5 Å as of 2023 [6]. Amongst other refinements of the technique, recent improvements in resolution are due to better cameras [7], improvements in the algorithms that reconstruct 3D molecular volumes from images [8– 11] and improvements to TEM software and hardware that allow semi-autonomous collection of larger numbers of particles [12, 13].

A recent advance in the field has been the use of beam shift – using the beam deflection coils in the upper objective lens system – to acquire from nearby regions of the grid instead of a slower mechanical shift of the microscope stage [13]. A compensating deflection of the beam in the lower objective lens system (known as image shift) shifts the beam back onto the camera, and modern microscope software uses stigmators to balance the coma aberrations induced by a beam tilt acquisition, with additional refinement and correction of the aberrations in SPA software [11]. When coupled with Köhler illumination (branded “Fringe-free illumination” by ThermoFisher Scientific and “Minimum Fringe” illumination by JEOL), which almost eliminates the Fresnel fringes typically seen at the edge of the illuminating beam, multiple images can be rapidly acquired per hole [13]. In this paper we demonstrate the use of square and rectangular apertures to generate corresponding square and rectangular beams in TEM so that even more images can be acquired on the same region of specimen. We show these aperture plates increase the number of images that can be recorded within the confines of a 1.2 *µ*m foil-hole of a cryo-EM grid by, on average, 75%, the precise value depending on the size of the acquisition field of view relative to the foil hole. We present designs for aperture plates that can be fabricated in a modern nanofabrication facility for a modest cost and can be installed in microscopes with only a single day of instrument downtime. Beyond re-centering the motorized aperture plates no further adjustment or realignment of the microscope is required. Finally we demonstrate that these modifications are compatible with high resolution cryo-TEM by recording data on samples of vitrified Equine apo-ferritin, a popular cryo-TEM standard sample, and solving the structure to sub-1.8 Å resolution.

### SQUARE AND RECTANGULAR APERTURES REPRESENT A AN APPROXIMATELY 50% MORE EFFICIENT USE OF SAMPLE AREA IN LOW-DOSE TEM

In cryo-TEM and other low dose modes the specimen will be damaged by too high an illuminating flux, typically around 40 e/Å^2^ for most bio-molecules [2], so to maximise the signal-to-noise ratio, each region is only exposed to the beam once. With a round beam expanded to fully cover the camera there will always be parts of the specimen exposed to the beam that are not acquired on the camera. In this section we calculate the extra efficiency of acquisitions, defined here as the fraction of cryo-EM foil grid that can actually be acquired assuming no area can be illuminated more than once, possible if a square or rectangular beam is used instead of a round beam.

Shown in Fig. 1 are cases where we have a (a) square or (b) rectangular (eg. the Gatan K3) camera and a round beam. The beam size is chosen such that it fully covers the camera and the acquisition efficiency *e* is the quotient of the area of the camera and the full area of the bea<u>m</u>. Assuming a perfectly uniform beam with diameter 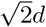 to fully cover a square camera with side length *d* this efficiency would be

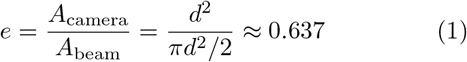

**FIG. 1.**
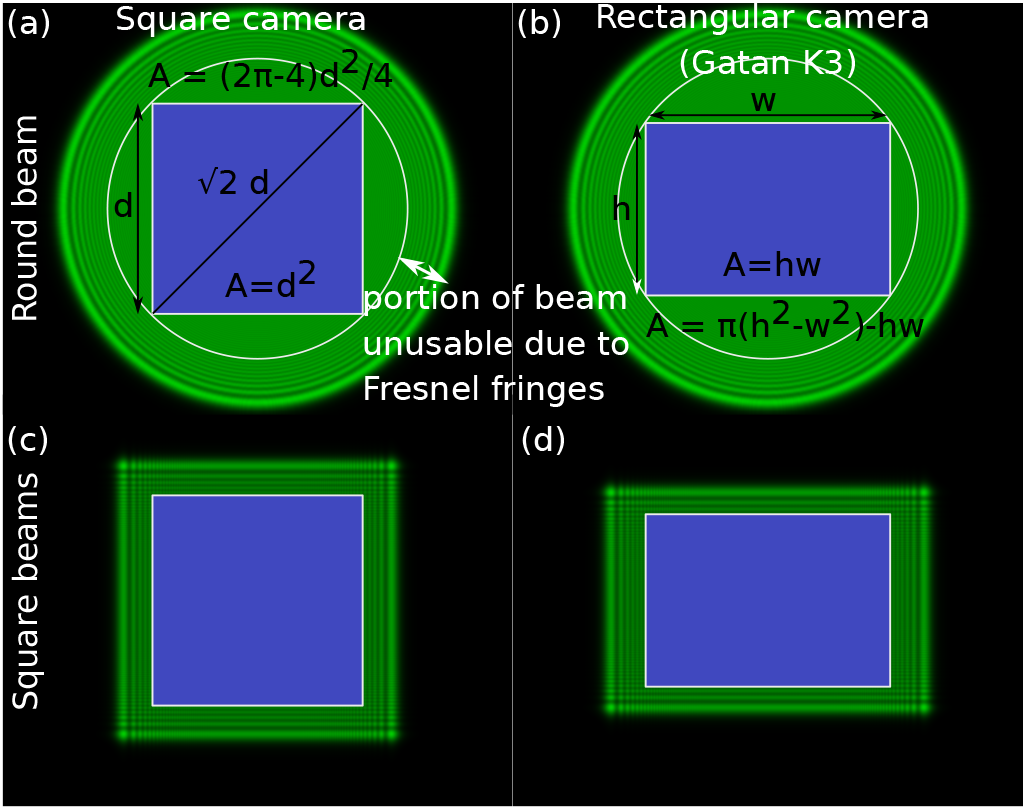
Square and rectangular beams result in a a much larger fraction of the illuminated sample area actually being acquired as shown in the schematics for a round beam and (a) square and (b) rectangular (eg. gatan K3) camera and for (c) a square beam and square camera and (d) a rectangular beam and rectangular camera.

Where a square or rectangular aperture is used, eg in Fig. 1 (c) and (d), the beam dimensions are much more sympathetic to the camera dimensions and the ratio of the area actually acquired to the area of specimen illuminated is improved. For a perfectly uniform square or rectangular beam that perfectly matches the underlying detector the efficiency defined in Eq. 1 would always be 1, in this simple case the improvement in efficiency in going from a round beam to a square beam would be 56%.

As in most TEMs, the beams in Fig. 1 all exhibit Fresnel fringes at the edges of the illumination that needs to be excluded from the images. Beam fringes result from the condensor aperture not being at the exact conjugate image plane to the specimen and means that the beam size always has to be larger than the camera size. We will briefly discuss the consequent impact on efficiency here. Fresnel fringes are markedly reduced on systems with modifications to address the issue - eg. Thermo Fisher refers to this as “Fringe-free illumination”, for example we measured a noticeable fringes extending 75 nm into our beam on our Thermo Fisher Arctica microscope but fringes only measuring 1 nm on our Thermo Fisher Krios G4 with the Fringe-free illumination feature. In appendix we show that square and rectangular apertures result in mild improvements for the ratio of specimen area recorded by the camera to the area illuminated by fringes. Having established this fact, to simplify discussion in the remainder of the section we consider the case of fringe-free illumination where the size of fringes are negligible.

We now discuss the acquistion area efficiency in the context of single particle cryo-TEM. In this technique, a grid containing a regular array of holes in a support film (usually carbon or gold) such as that sold by Quantifoil or the “C-flat” grid manufactured by Protochips is preferred. The consistency of hole size and their regular placement permits automated acquisition of the ice region inside the hole by software on the TEM such as serial-EM or EPU. Advances in TEM software packages mean that schemes where there are multiple acquisitions per foil-hole, colloquially referred to as “multi-shot” acquisition, are now common. In Fig. 2 we compare the efficiency of multi-shot acquisitions for different pixel sizes and different cameras and beam types. We assume that the areas of specimen illuminated in successive acquisitions cannot overlap (ie. no region of the specimen can be exposed to the beam more than once, even if that region is not actually acquired on) and that no support film in the field of view is permitted. Fig. 2 plots the fraction of the hole that is acquired using a round beam with a 4096 x 4096 pixel camera (modelled on the ThermoFisher Falcon IV installed on our Titan Krios) and a rectangular 5760*×* 4096 pixel camera (modelled on the Gatan K3 camera insatlled on the Krios microscope). Multi-shot configurations were drawn from the most efficient circle within circle packing schemes described in Ref. [14] for round beams and from Ref. [15] for square beams within a circle. For the case of a 5760 *×* 4092 pixel K3 camera we developed our own tiling schemes, all of which are plotted in Fig S1 of the supplementary. The efficiency, or fraction of the foil hole that is actually imaged, is plotted for each configuration and the number of acquisitions in a foil hole is annotated for a few examples atop the plot. Example configuration schematics are inset in the plot for the K3, Falcon and square beam examples at a pixel size of 0.65. For a pixel size of less than 1.0, that which is commonly used in cryo-EM, an average fraction of 0.38 and 0.31 of a foil-hole is covered by the Falcon and K3 with a round beam respectively. For the Falcon IV the fraction a square beam configuration can achieve is 0.64 for these conditions, approximately double that of the round beam case. Taken over the plotted range, a pixel size varying from 1.5 Å to 0.4 Å the average improvement was calculated to be 73% for the square Falcon IV and 74% for the rectangular K3 camera.

**FIG. 2.**
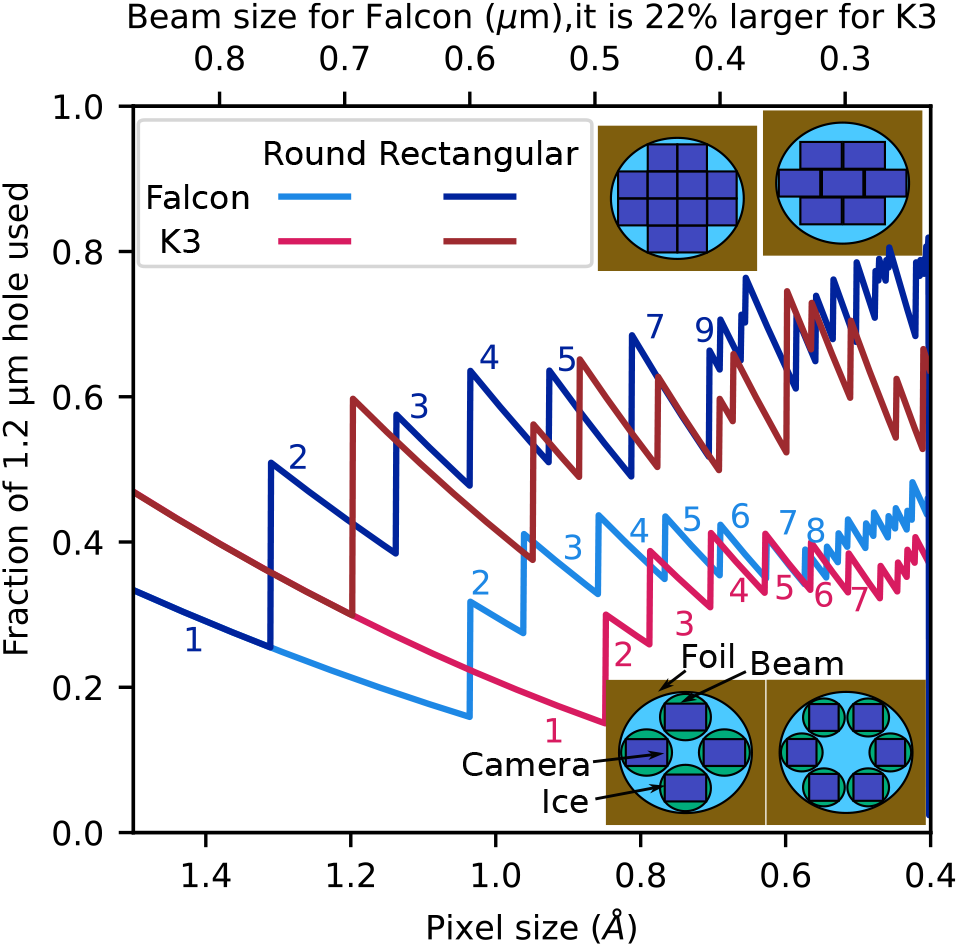
In single particle cryo-TEM images are acquired in an ice-covered foil-hole. Depending on the configuration of beam and camera (round, square or rectangular) a different fraction of the hole would actually be imaged. Plotted in is the total fraction of the foil hole that can be *both* illuminated and acquired using a square and round beams with a square (eg. Thermo Fisher Falcon IV) or rectangular (eg. Gatan K3) camera if no foil is tolerated in the images as a function of pixel size (Å). The number of acquisitions that can be fitted into a multishot configuration is annotated on the figure for example cases, next to the plotted lines. Schematics of different example multishot configurations for each of the camera beam combinations at a pixel size of 0.65 Å are also inset. A square or rectangular beam maximises the number of non-overlapping acquisitions that can fit into one of these foil holes and thus the total fraction by approximately 73% on average.

## DESIGN AND FABRICATION OF APERTURE PLATES FOR A THERMOFISHER TEM

In this section we describe the design and fabrication of square and rectangular aperture plates for our ThermoFisher Arctica G2 and Krios G4 instruments. We describe a process for producing affordable aperture plates that provide appropriately sized beams for different magnifications and cameras within a single plate that, once loaded to the microscope, require minimal instrument realignment and downtime. Modern ThermoFisher TEMs feature exchangeable aperture plates with a spring-loaded rod holding up to 3 plates, each 3.04 mm in diameter, See Fig. 3(a). Instruments buit by JEOL, an alternative TEM manufacturer to ThermoFisher, feature a similar system for interchanging aperture plates though the diameter of the plate can vary between 2 and 3 mm depending on the model of microscope, so these designs should be adapatable to these microscopes though fabrication of plates for these microscopes is not something we’ve attempted yet. To load an aperture the rod is pulled back and the plate is dropped into the circular cut-out at the end of this groove, as shown in Fig. 3(b). The aperture can then slide into position inside the groove and once the rod is released the tension of the spring will hold the apertures plates in place. Typically only one aperture is present at the centre of these plates. To anticipate the fact that electron microscope lenses rotate the electron beam as the strength of the condenser lens system is adjusted to form different sized beams (this is of course not noticeable with a circular beam), we took the approach of offering multiple rotated copies of each desired aperture on a single plate so that there would be an beam approximately oriented to the camera at each magnification. This approach was modelled on that of Zeltmann et al. [16] who also milled multiple aperture holes within the same plate.

**FIG. 3.**
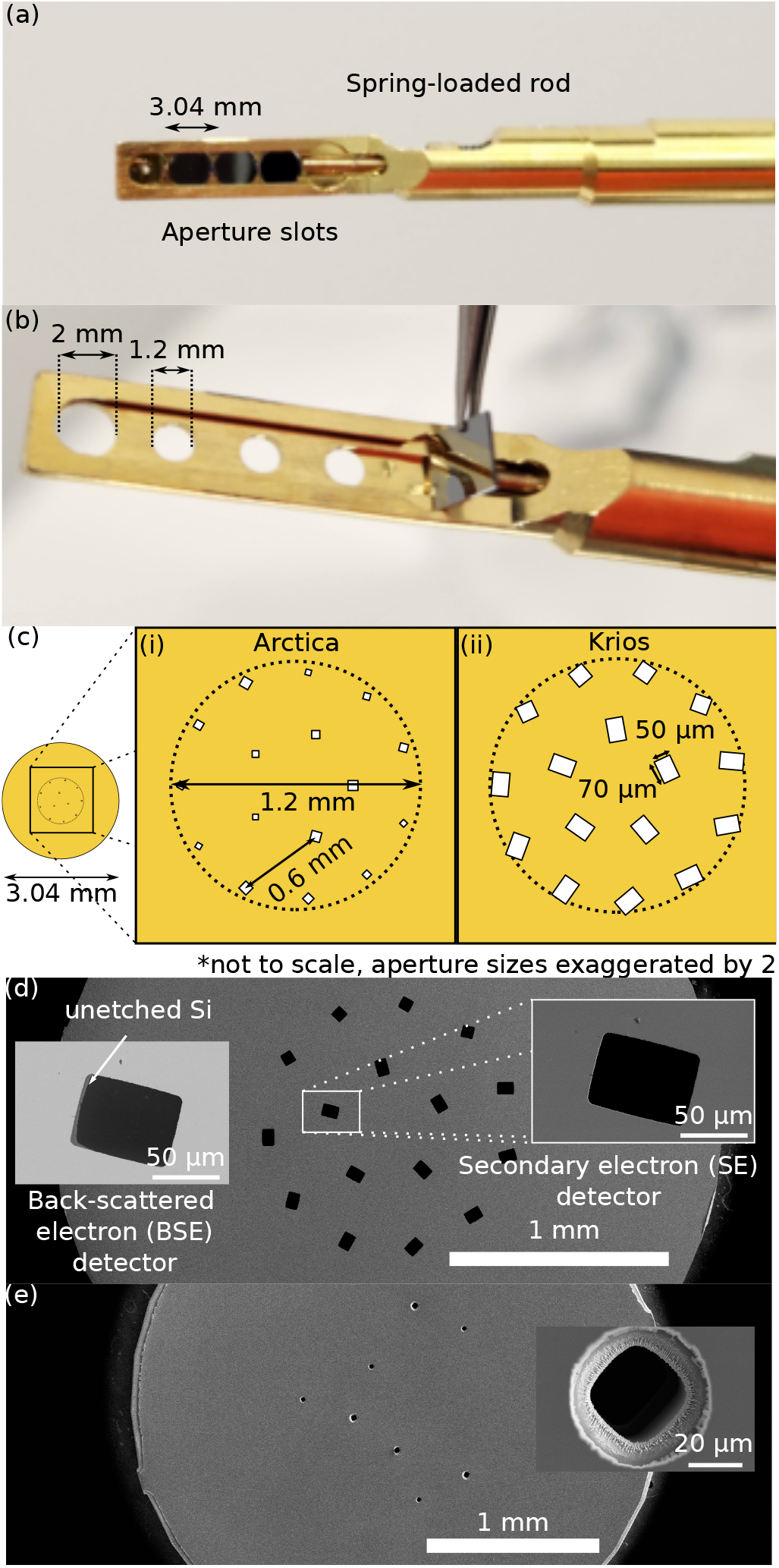
(a) A ThermoFisher aperture rod with four aperture plates loaded and (b) with the rod pulled back. The 2 and 1.2 mm holes determine how the space available on the plate for fabricating apertures. Shown in (c) are schematics of the designs used in fabrication with SEM images of the final aperture plates for the (d) Krios and (e) Arctica. Only the larger holes in the Arctica plate successfully milled.

### Arctica G2 aperture plate

The Arctica is a 2-condenser lens microscope which means that the beam is only parallel at one single setting of the second condenser (C2) lens and only one size of parallel beam can be formed for a given size of C2 aperture. In our aperture design, as well as offering square apertures, we additionally aimed to improve the flexibility of these instruments by giving more aperture options that could be used to give beams that better matched the camera field of view at the different magnifications used for single particle cryo-TEM. This should improve the flexibility of these microscopes for multi-shot cryo-TEM acquisitions.

The area of the 3.04 mm plate that can be used for aperture fabrication is limited by holes on the underside of the rod measuring approximately 2.0 mm for the slot closest to the tip of the rod and 1.2 mm for the three slots further down the rod as shown in Fig. 3(b). To determine how closely the apertures needed to be packed within a single plate we estimated the size of the beam at the C2 aperture plane, this was achieved by inserting the 20 *µ*m C2 aperture and translating it from its usual position at the microscope’s optic centre until the beam disappeared (ie. the C2 aperture moved past the edge of the beam). The point at which this occurred was found to vary with spot size (consistent with the fact that changing spot size involves coupled changes in the C1 and C2 condenser lens strength to magnify and de-magnify the electron spot at the gun plane) but the largest beam radius at the C2 plane was found to be 300 *µ*m for the Arctica. We took this as the desired separation between different apertures on a single plate.

We created two designs, one where all the aperture holes were located inside a 1.2 mm disc and another where all were located inside a 2 mm disc, the latter design provided more space for the apertures but less options for slots in the aperture rod, see Fig. 3(b). As mentioned previously, we wanted the ability to create square beams at each of the four magnifications we commonly used for single-particle cryo-TEM, each detailed in Table I, and we created four of each 15^*°*^ aperture rotation for a total of 4*×* 4 = 16 apertures. We measured that the fringes at the edge of the illumination and average beam position drift over the course of an acquisition to both be approximately 75 nm so added 300 nm to the sensor size (i.e. an extra 150 nm margin to each side of the beam) to calculate the desired size of the square beam. To determine the physical aperture hole size we measured the multiplicative factor that related aperture to beam size at the parallel condition to be 29.4 using the 20 *µ*m aperture already installed on the Arctica and used this factor to calculate the aperture hole size that would result in a desired beam size. A schematic of the final design is shown in Fig. 3(c)(i), the optimal layout of the 16 aperture holes to ensure maximum separation between different holes whilst still fitting inside a 1.2 mm circle is taken from [14], which ensured that apertures where spaced just over 300 *µ*m apart.

**TABLE I.** Different acquisition magnifications commonly used in our Talos Arctica microscope with the calibrated pixel size, size of the acquisition region on a 4096*×* 4096 pixel K2 camera, the required size of the beam, factoring in the 75 nm Fresnel fringes and the final size of the aperture which factored in another 75 nm to account for beam drift during an acquisition.

| Magnification | Pixel size<br>(Å) | Sensor size<br>(nm) | Beam size<br>(nm) | Aperture size<br>($\mu\text{m}$) |
| --- | --- | --- | --- | --- |
| 79,000 | 1.71 | 656 | 956 | 29.5 |
| 100,000 | 1.33 | 510 | 820 | 24.5 |
| 130,000 | 1.03 | 395 | 695 | 20.5 |
| 165,000 | 0.788 | 302 | 602 | 17.5 |

### Krios G4 aperture plate

Our ThermoFisher Krios G4 is a 3 condenser lens microscope and, in TEM mode, the second and third condenser lens (C2 and C3) strengths are software linked to ensure that a parallel beam can be formed for a range of beam sizes so only a single size of aperture needed to be made for every rotation. Since this TEM is equipped with both a K3 (4096 *×* 5960 pixel) and Falcon (4096*×* 4096 pixel) camera, both square and rectangular apertures needed to be fabricated into the same plate. The initial design, shown in Fig. 3(c)(ii) featured square and rectangular apertures that measured 50 *µ*m in the smallest dimension, with 4*×* 15^*°*^ rotations covering a 60^*c*^*irc* range for the square apertures and 12*×* 15^*°*^ rotations for the rectangular apertures. Upon loading this plate to the microscope we realised that this size of aperture was insufficient for forming a small enough parallel beam to match the camera at higher magnifications than 130,000. Attempting to reduce the beam size further meant that microscope beam mode changed from “parallel” to “condensing”, resulting in a non-uniform beam brightness that was not usable for acquiring high-quality data. We have since revised the design to feature apertures that measure 30 *µ*m in the smallest dimension so that sufficiently small beam sizes to cover the full range of magnifications are achievable.

#### Fabrication of aperture plates by reactive ion etching (RIE)

Aperture designs, specifically the positioning of the vertices of each aperture on the 3.04 mm plate were generated using a Python script included in the supplementary information. The ezdxf Python library [17] was used to write these vertices as Polygon objects in the AutoCAD drawing exchange format (dxf) and these are also included in the supplementary material. The dxf designs were transferred to a Chrome / Glass photo-mask that is used in UV photolithography as a 1:1 transfer mechanism.

200um thick silicon was chosen as the substrate of choice for the apertures due to silicon’s relative strength and its ability to be etched into shapes with high aspect ratios that were necessary for the apertures. To transfer the pattern from the photo-mask to the Silicon wafer, AZ40XT photo-resist was spin coated and soft baked onto the Si substrate at about 25um thickness. The pattern transfer was then achieved by 365nm UV exposure of the patterned photomask onto the photo-resist. Subsequent development of the pattern removed the UV exposed areas of the photo-resist, creating areas on the Silicon that could then be removed by reactive ion etching (RIE) using a Oxford PlasmaLab RIE machine. Areas not targeted for etching were protected by the remaining photo-resist. After the features were formed into the Silicon substrate, the remaining sacrificial protective resist was removed using RIE with O_2_ plasma. To minimise charging of the apertures which would likely produce strong probe aberrations, 1um of conducting Au film was then sputter deposited onto the Silicon wafer covering the apertures and making them conductive. Finally, the wafer was cut into individual pieces correctly sized to fit into the Transmission Electron Microscope’s aperture holding mechanism.

In this single fabrication run 32 aperture plates were made, demonstrating a process that can easily scale to producing aperture plates for many different microscopes at a cost that is minimal in comparison to usual component costs for electron microscopes. Two example aperture plates were then inspected in a ThermoFisher Teneo Volumescope SEM, and these images are shown in Fig. 3 (d) for the Krios design, showing the top (beam-facing, gold plated) side and (e) for the Arctica design, showing the bottom (opposite to beam, unplated) Si side. For the Krios design, Fig. 3 (d), all holes milled successfully though there is some rounding of the vertices of the aperture, this is not overly problematic because this would impact only the very edge of the beam which is rendered unusable by Fresnel fringes anyway. Also visible was the appearance of a small rim of silicon that is more easily visible in the back-scattered electron (BSE) detector SEM image of Fig. 3 (d) where it is marked with a white arrow. Both of these defects had a mild impact on the performance of the apertures since the match with camera was not as good as would have been the case if the original designs were etched perfectly. Nonetheless overall quality of the apertures was pleasing and exceeded expectations for a prototype fabrication run. For the Arctica aperture plates only the larger 24.5 *µ*m and 29.5 *µ*m apertures milled successfully, these being at the lower end of what is achievable with reactive ion etching of 200 *µ*m thick Si wafers. Before loading into the condenser aperture holder of the microscope the apertures a gaseous N_2_ jet was used to remove dust and the plates were plasma cleaned for 5 minutes in an O/Ar Fischione 1020 plasma cleaner. The ThermoFisher Krios G4 microscope column was vented and the existing 70 *µ*m aperture plate was replaced with the design shown in Fig. 2 (c), normal vacuum was achieved after an overnight pumping cycle.

#### Data collection using square and rectangular beams in cryo-EM

Once the new apertures were loaded into the ThermoFisher Krios G4 their positions were found and then recalled by adjusting the x and y positioning of the aperture until the apertures came into the field of view. The ThermoFisher utility `tadBhvApertures.exe`, which is used for engineer configuration and alignment of the apertures, was particularly useful because it gave aperture plate positioning values that could be manually recorded for each aperture opening and then used to quickly bring a chosen aperture back to position. As discussed in earlier sections, each magnification implies a different camera field of view and therefore a different size of beam, modifying the size of the beam by adjusting the condenser lenses results in some rotation of the beam. We found that generally the same aperture gave a result within 10^*°*^ of the camera orientation for most of the magnification ranges used for cryo-EM acquisition and a different aperture only being necessary for the lower magnifications (*<* 100, 000 *×*). An image of the beam with one of the rectangular apertures inserted, condensed to be just smaller than the camera is shown in Fig. 4, the effects of the aperture being not perfectly rectangular square due to incomplete etching of some Si, as shown in the SEM images in Fig. 3 are visible in the final beam. Because this instrument is a ThermoFisher Krios G4 with fringe-free or Köhler illumination the fringes are approximately 1 nm in size, as can be seen in Fig. 4(a), whereas they were measured to be 75 nm on our Arctica microscope. An example multi-shot setup from the ThermoFisher EPU single particle Cryo-EM automated acquisition software ins shown in Fig. 3(b) this time for the square Falcon apertures, burning the surrounding carbon with the beam was found to be an accurate method of giving a good template to help guide setup of the multishot pattern.

**FIG. 4.**
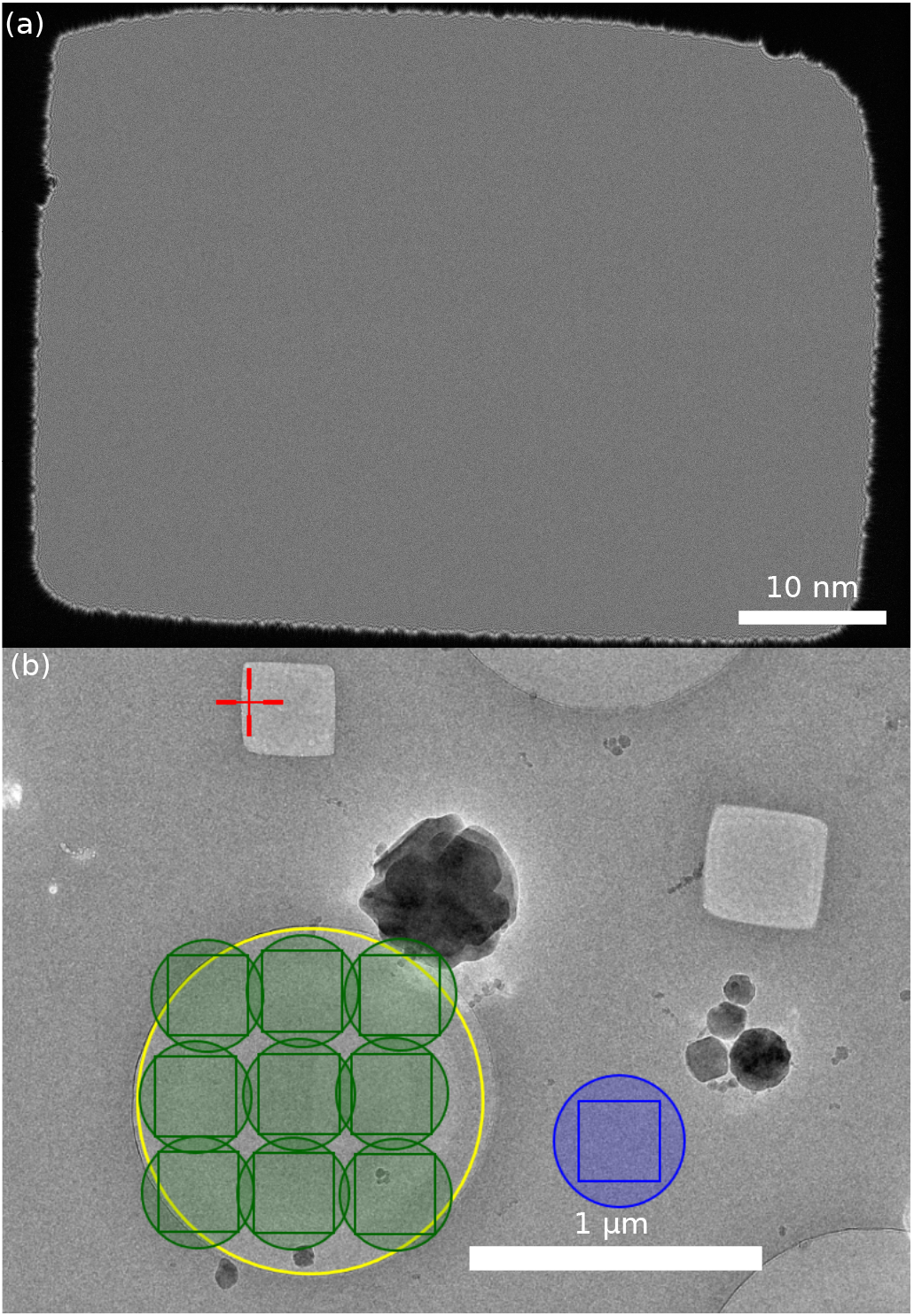
(a) Image of the beam with a rectangular aperture recorded on the K3 camera. alignment is to within 10^*°*^ with the camera and fringes at the edge measure less than 1 nm with the fring-free illumination system. Imperfect etching seen in Fig. 3(b) is visible in the beam resulting in rounded edges. (b) Set up of a multishot acquisition on a carbon grid in the EPU single particle cryo-EM software, this time for the square beam. The ice is purposely burnt by the beam to create a visual template of the square beam to assist in optimal design of a multishot pattern.

To benchmark cryo-EM acquisition with the new aperture design and demonstrate their potential for high-resolution single particle analysis we recorded datasets with the new apertures the ThermoFisher Falcon IV camera and square aperture and Gatan K3 and rectangular aperture combinations. We also present data recorded using the same cameras with the standard 50 *µ*m round aperture as a comparison dataset. We used Sigma-Aldrich equine apoferritin (product number A3641), 4 *µ*L of the solution was pipetted onto gold 1.2/1.3 Quantifoil Aultrafoil grids, which were glow discharged for 180 s in Quorum Glocube with a 15 mA current. The grids were prepared using a ThermoFisher Vitrobot Mk IV, set to room temperature (22^*°*^) with a humidity of 95%, using a blot force of -1 and blot time of 4 s before plunging into liquid ethane. Key parameters and metrics of the datasets are shown in the table of Fig. 5. The ratio of movies to final particles varies between the datasets mainly due to observed particle densities in the different preparations. The final number of particles was similar enough to play a minor role in the final resolution. For each dataset the processing was as follows, movies were aligned using the patch motion correction algorithm of the Cryosparc cryo-EM processing software [10] with CTF-fitting also performed using the patch motion correction algorithm and particle picking was performed using a template generated by 2D-classification of 100 manual picks. A subset of quality particles were selected through two rounds of 2D-classification and the particles then went into 3D reconstruction, the initial model being generated by Cryosparc’s Ab-initio model routine. Particles were split into different optics groups using the EPU_group_ AFIS.py python script [18], which parses metadata .xml files outputted by Thermo Fisher’s EPU software to extract beam-tilt values for each acquisition and clusters together acquisitions, this grouping was then imported to Cryosparc using an in-house script (get_exp_group_id. py, which is included in the supplementary materials). 3D refinement in Cryosparc with per-particle defocus refinement and per-group CTF refinement (including beam tilt and shift, trefoil and tetrafoil aberrations) produced a maps with resolutions between 1.76 Å and 1.89 Å resolutions as measured using the 0.143 Fourier shell correlation (FSC) criteria [4], suggesting that square or rectangular apertures did not hinder the microscope’s ability to reach high resolutions.

**FIG. 5.**
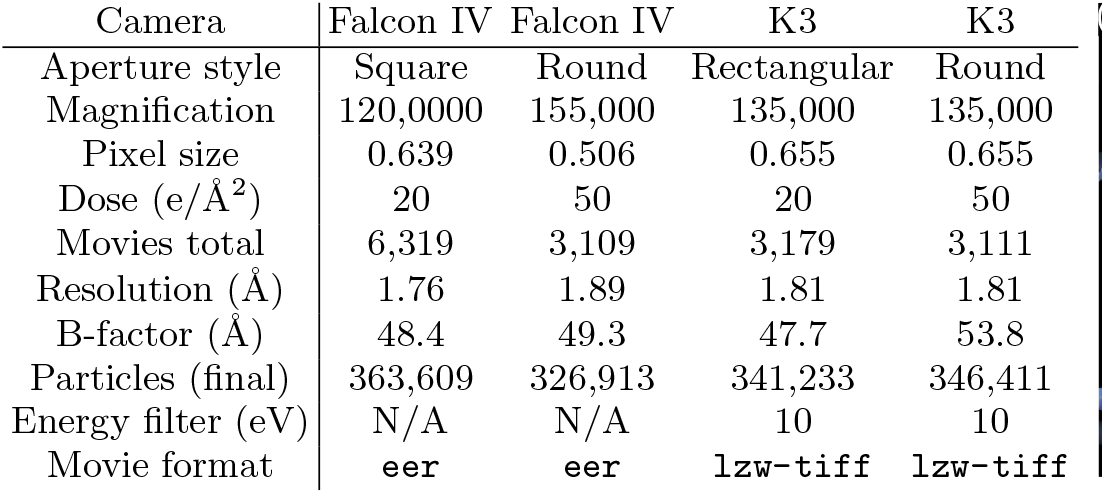
Parameters and speed and resolution metrics for the acquisitions presented in the manuscript

A slice of the structure, visualized using ChimeraX [20], emphasising the visibility of aromatic carbon rings in phenylalanine and tyrosine amino acid side chains for the Falcon IV dataset with a square aperture is shown in Fig. 6(a) overlaid with a stick molecular model from pdb entry 6rjh [19]. A complete view of all the models for each dataset, also generated using ChimeraX [20], is shown in Fig. 6(b) with all the FSC curves in a single plot displayed in Fig. 6(c) with the 0.143 FSC criterion indicated. Separate CTF refinement including spherical aberration (C_*S*_) in Cryosparc did not indicate that the aperture plates were inducing extra aberrations in the microscope’s objective lens system.

**FIG. 6.**
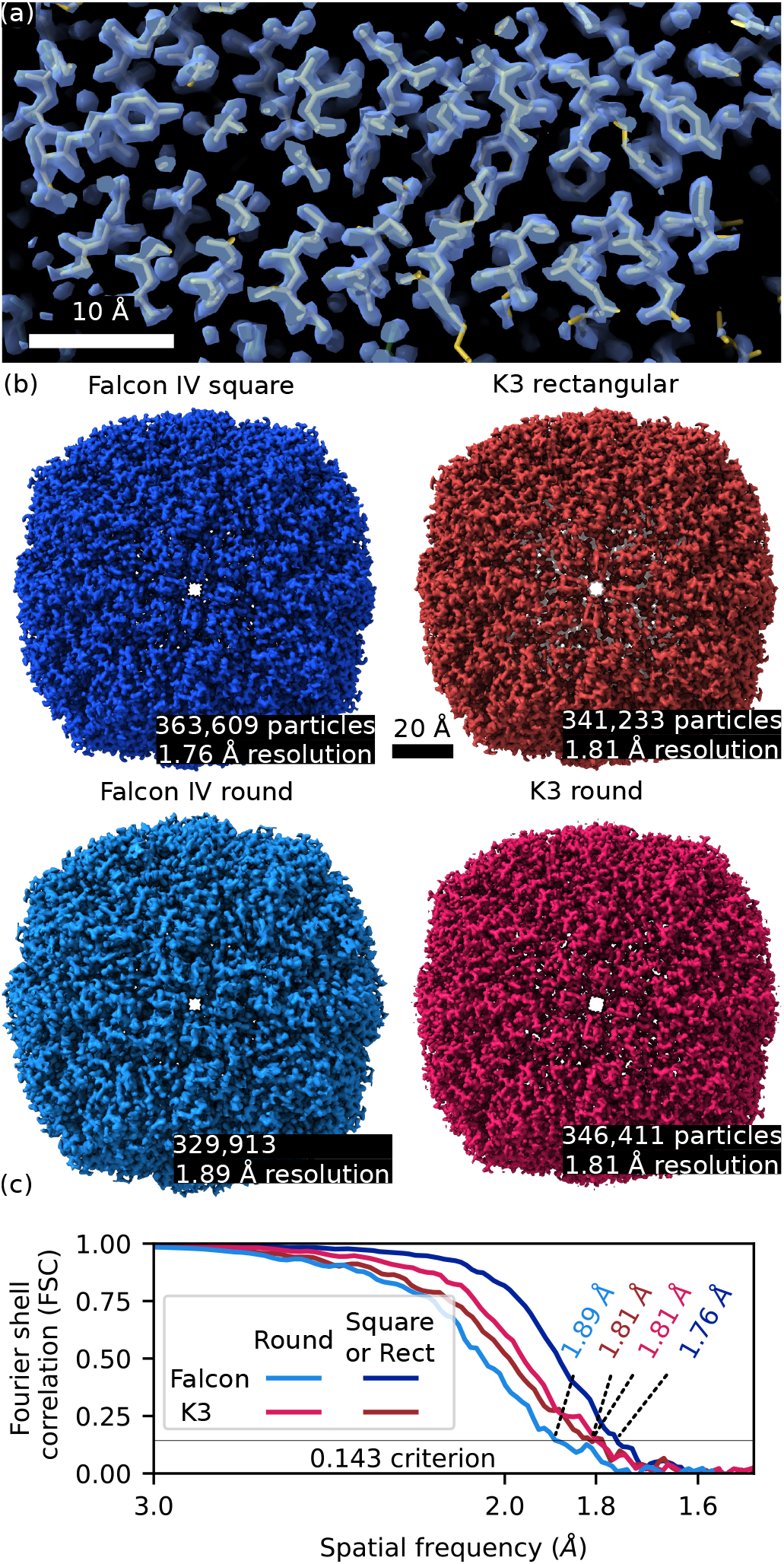
Reconstructed models of Apo-ferritin. Shown in (a) is a cross section of the reconstructed map with pdb entry 6rjh [19] overlaid, aromatic rings from phenylalanine and tryptophan amino acid side chains are clearly resolved. (b) Reconstructed volume visualized in Chimera X [20]. (c) Fourier shell correlation (FSC) plots to gauge resolution at the 0.143 criteria [4].

## CONCLUSION

In the acquisition schemes presented here square and rectangular beams lead to an improvement in the fraction of the grid that could be acquired by approximately 75 %. We have presented and fabricated designs for square and rectangular aperture plates that produce corresponding square and rectangular beams. A demonstration dataset resulted in a final map with a resolutions at or below 1.8 Å demonstrating that these aperture plates are appropriate for high resolution data collection.

## ACKNOWLEDGEMENTS

We acknowledge use of the ThermoFisher Krios G4 cryo-TEM and ThermoFisher Teneo VolumeScope SEM at the Ian Holmes Imaging Centre of the Bio21 Institute at the University of Melbourne. Mr. Philip Francis is thanked for training and assistance with the ThermoFisher Teneo VolumeScope SEM. This work was performed in part at the Melbourne Centre for Nanofabrication (MCN) in the Victorian Node of the Australian national Fabrication Facility.

In the late stages of manuscript preparation we became aware of the work of Chua et al. [21] which was recently published on the BioR*χ*iv and is similar to the work presented here.

The next generation of aperture plates are being fabricated by Norcada (https://www.norcada.com/), a MEMS and photonics fabrication facility in Edmonton, Alberta, Canada. We also thank them for initial discussions on aperture fabrication in the early stages of the project.

Dr. Stephen Zeltmann, Dr. Jim Ciston and Dr. Colin Ophus from the National Centre for Electron Microscopy (NCEM) at Lawrence Berkeley National Laboratory, California, USA are thanked for initial discussions on fabrication of condenser aperture plates for ThermoFisher Titan TEMs and the multiple apertures in a single plate design was largely inspired by their example.

All in-house software used in this publication including the script to generate DXF design files is available in the supplementary materials.

## Appendix

### Fresnel fringes in square and rectangular apertures

In this appendix we consider the area of the beam that is unusable due to Fresnel fringes, assumed to be of constant width *f*, for each of the cases considered in Fig. 1, we show that square and rectangular beams result in the same fraction of area of the beam lost to fringe thus still provide a benefit even in cases were fringe width is significant. In most cases we can assume that the radius of the uniform area of the beam *d* are much larger than the fringe width *f* (ie. *d ≫ f*). For a square ca<u>m</u>era with side length *d* and round beam with radius 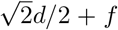, as s<u>h</u>own in Fig. 1(a), the total beam area will be *A* = *π* (2*d/*2 + *f*)^2^ and the ratio of the area of the fringes to the area of the camera will be:

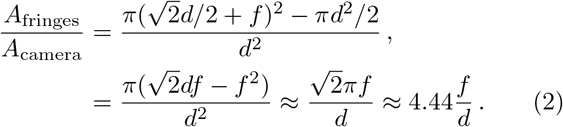

We now consider the same square camera but now with a square beam, ie. the case shown in Fig. 1(c). We assume a square beam and camera with side lengths *d* + 2*f* and *d* respectively and the ratio of the area of the fringes to the area of the field of view of the camera will be:

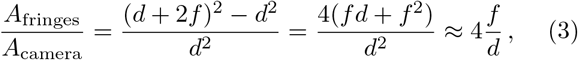

which is slightly smaller than for the round case in Eq.(2), meaning square beams still provide a modest advantage in the case of large Fresnel fringes at the edge of the beam.

For the case shown in Fig. 1(b), a rectangular camera with longer side a factor o<u>f</u> 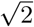 times its shorter side *d*, the beam radius will be 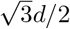 to cover the c<u>a</u>mera and the total area of the beam will be 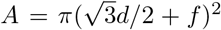 and the ratio of the area of the fringes to the area of the field of view of the camera will be

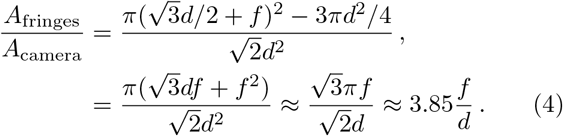

For the equivalent case of a rectangular beam *and* camera,

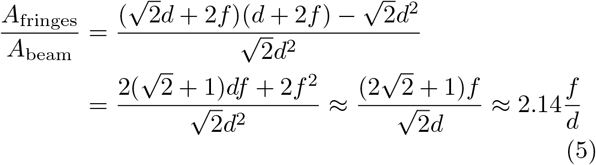

similar to the square camera case, this ratio is smaller than for was the case in Eq. (2), meaning rectangular beams still provide a modest advantage in the case of large Fresnel fringes at the edge of the beam.

