## Supplementary for "Fitting a square beam in a square camera: novel condenser apertures for low-dose transmission electron microscopy" (from another version of this preprint: version 1, 2023-08-15)

August 14, 2023

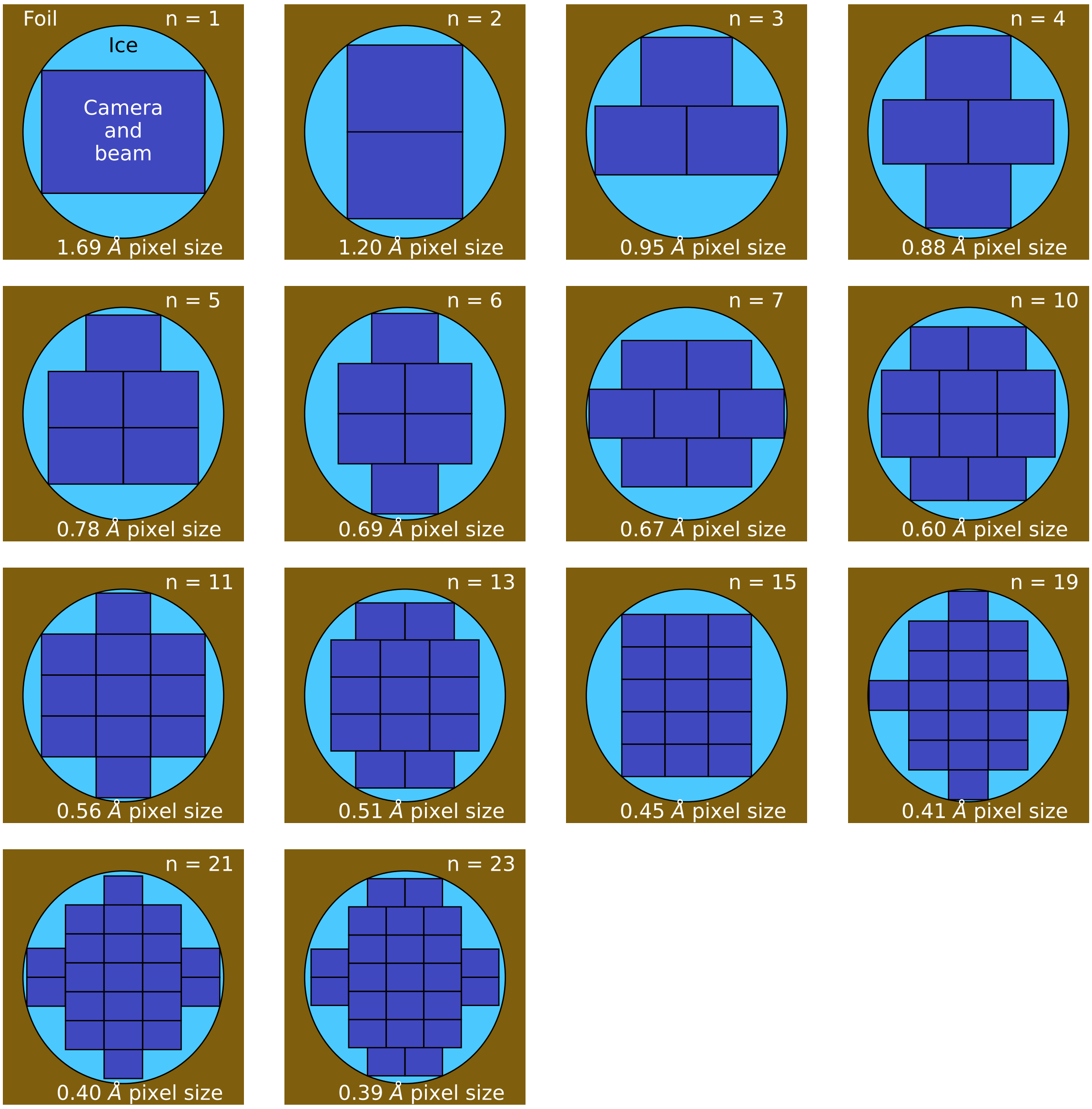


Figure S1: Multishot configurations for a rectangular K3 camera and matching beam ($5760 \times 4092$ pixel) camera inside a 1.2 $\mu$m diameter foil hole used to generate the efficiency measurements presented in Fig. 2 of the main text.
